# Hybrid Ornstein–Uhlenbeck–Branching Modeling of Pediatric Leukemia Evolution: A Computational Extension and Cohort-Level Application

**DOI:** 10.1101/2025.11.07.686044

**Authors:** Seung-Hwan Kim

## Abstract

Pediatric cancers evolve under developmental constraints that limit mutational diversity yet preserve adaptive potential. A computational extension of the Hybrid Ornstein–Uhlenbeck (OU)–Branching framework was developed to model clonal diversification and phenotypic stabilization in pediatric leukemia. The OU component captures mean-reverting dynamics representing developmental homeostasis, while the branching component introduces stochastic lineage bifurcation and extinction. Using de-identified clinical metadata from *Ahlgren et al*. (*Nature Communications*, 2025; 16:8964), the model simulates patient-specific evolutionary trajectories across relapse categories and disease subtypes (B-ALL, T-ALL, MPAL, AML). Simulations reproduce observed clinical trends—rapid relapse and limited diversification in early or refractory KMT2A-r ALL, and slower, therapy-resistant relapses in AML. Group- and patient-level analyses demonstrate how the balance between stabilizing selection (*θ*) and diversification rate (*λ)* determines clonal persistence and phenotypic drift. This computational implementation provides a quantitative framework for linking developmental constraint, clonal diversity, and therapeutic response in pediatric malignancies and establishes a tractable platform for model-driven hypothesis testing and translational oncology.

**Statement of Relationship to Prior Work:** This preprint represents a computational and cohort-level extension of the hybrid OU–Branching framework introduced in the author’s manuscript currently under review at Frontiers in Oncology (“A hybrid Ornstein–Uhlenbeck–branching framework unifies microbial and pediatric tumor evolution,” Manuscript ID 1727973). The Frontiers paper focuses on experimental validation and cross-domain analogies between microbial long-term evolution experiments (LTEE) and pediatric tumor evolution, emphasizing biological interpretation. In contrast, this preprint focuses on clinical modeling, patient specific simulations, and computational methods applied to pediatric KMT2A-rearranged leukemia. No data, text, or figures are duplicated from the in-review article. All code and simulations presented here are novel and will be released upon publication.

## 1. Introduction

Classical models of cancer evolution have relied on Markovian or birth–death processes, which assume memoryless transitions among discrete mutational states. Rooted in mid-20th-century stochastic population theory (Kendall, 1948; Athreya & Ney, 1972), these frameworks have long been applied to describe tumor growth, mutation accumulation, and clonal expansion. Multi-type branching processes and continuous-time Markov chains have yielded analytical insights into mutation waiting times, driver– passenger dynamics, and therapy resistance (Iwasa et al., 2006; Beerenwinkel et al., 2007; Bozic et al., 2010; Durrett, 2013; Michor et al., 2004), effectively capturing adult tumor progression where mutation rates and fitness differences can be approximated as stationary and independent (Komarova & Wodarz, 2005). Yet these memoryless frameworks neglect adaptive feedback and developmental constraints, which are central to pediatric malignancies.

Pediatric tumors emerge within developmental contexts characterized by high cellular plasticity, low mutational burden, and strong stabilizing selection (Groebner et al., 2018; Ma et al., 2018; Huether et al., 2014; Mack et al., 2017). Originating from progenitor or embryonic lineages, their evolution is shaped by canalization, epigenetic regulation, and non-Markovian feedback loops that constrain accessible trajectories (Huang, 2011; Karin & Alon, 2021; Basanta & Anderson, 2013). Consequently, models that treat lineage evolution as independent of past states fail to capture the developmental and adaptive coupling intrinsic to pediatric cancers. While extensions of classical frameworks have incorporated feedback and plasticity through fitness landscapes or phenotypic-state transitions (Archetti et al., 2015; Lande, 1976; Hansen, 1997; Butler & King, 2004; Uyeda et al., 2011; Hansen & Martins, 1996), and evolutionary quantitative genetics has employed Ornstein–Uhlenbeck (OU) processes to describe trait evolution under stabilizing selection (Gröbner & Pfister, 2020), to our knowledge, no existing model unifies adaptive constraint with branching diversification in a biologically grounded cancer framework.

To address this gap, a **Hybrid Ornstein–Uhlenbeck (OU)–Branching model** was formulated to integrate adaptive mean reversion and lineage diversification. The OU component captures stabilizing selection toward an equilibrium trait value (*μ*) through selection strength (*θ*) and diffusion variance (*σ*^2^). In contrast, the branching component represents clonal diversification at a rate *λ*, reflecting mutational or epigenetic bifurcation events. This dual structure provides a mechanistic basis for describing how developmental constraint and evolutionary exploration jointly govern clonal persistence, extinction, and phenotypic drift in pediatric cancers.

Applied to the KMT2A-rearranged leukemia cohort reported by Ahlgren et al. (Nature Communications, 2025; 16: 8964), the Hybrid OU–Branching framework reproduces observed patterns of rapid relapse in early-stage acute lymphoblastic leukemia and slower, therapy-resistant recurrence in acute myeloid leukemia. Through this integration of adaptive feedback and stochastic diversification, the model establishes a tractable foundation for quantifying evolutionary constraint and variability in pediatric malignancies.

## 2. Methods

### 2.1 Mathematical Foundations

The Ornstein–Uhlenbeck (OU) process describes the temporal evolution of a continuous trait under stabilizing selection and stochastic drift (Øksendal, 2013; Hansen, 1997; Lande, 1976). The stochastic differential equation is defined as

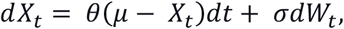

where *θ* > 0 denotes the selection strength toward the equilibrium trait *μ, σ*is the diffusion coefficient, and *W*_*t*_ represents a Wiener process. The deterministic drift term *θ*(*μ* − *X*_*t*_) generates mean reversion toward *μ*, while the stochastic term introduces random variability. At stationarity, the process converges to a normal distribution centered at *μ* with variance *σ*^2^/(2*θ*), reflecting stabilizing selection around a developmental optimum.

### 2.2 Lineage Diversification

To model clonal diversification, a branching component was incorporated into the OU framework. Each lineage follows OU dynamics and, with rate λ > 0, divides into two descendants inheriting parental traits plus Gaussian perturbations (Ethier & Kurtz, 1986; Durrett, 2013; Butler & King, 2004):

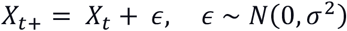

The expected number of lineages increases exponentially with rate λ, while the OU drift constrains trait variance, maintaining phenotypic stability amid diversification. This coupling allows the model to represent clonal expansion, extinction, and bounded diversification under stabilizing selection.

### 2.3 Infinitesimal Generator of the Hybrid OU–Branching Process

The infinitesimal generator unifies drift, diffusion, and branching in a single operator. For a smooth test function *f*(*x*), the generator is given by

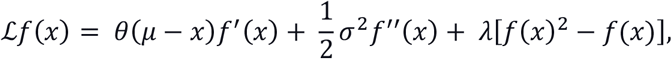

representing the simultaneous contribution of adaptive mean reversion, stochastic variability, and lineage bifurcation (Øksendal, 2013; Ethier & Kurtz, 1986). This formulation defines a continuous–discrete hybrid process capable of describing both gradual phenotypic evolution and discrete diversification events within tumor lineages.

### 2.4 Computational Framework

Hybrid OU–Branching, Brownian, and Markov processes were simulated in Python 3.11 using the Euler– Maruyama integration scheme for the continuous component. Branching was implemented as a Poisson process with rate λ, corresponding to the expected number of bifurcations per unit time, and a branching probability of *p*_*b*_ = 0.5 per lineage per event. All simulations employed fixed parameters (Supplementary Table S1), as the objective was to characterize model behavior rather than parameter estimation. Parallelized numerical solvers built on NumPy enabled the efficient simulation of thousands of lineages with adaptive time steps (*Δt*= 10^−3^) to ensure numerical stability and computational efficiency.

### 2.5 Clinical Dataset and Cohort Simulation

Clinical metadata were obtained from Ahlgren et al. (Nature Communications, 2025; 16:8964), specifically from SuppData1_ClinicalData. The cohort comprised 36 pediatric patients—25 diagnosed with B-cell, T-cell, or mixed-phenotype acute lymphoblastic leukemia (B-ALL, T-ALL, MPAL) and 11 with acute myeloid leukemia (AML)—encompassing 257 longitudinal samples collected at diagnosis, remission, and relapse. De-identified patient identifiers, disease types, and relapse groups (very-early, early, late, and remission) were extracted. Median relapse intervals (419 and ∼289 days for ALL and AML, respectively) were scaled by group-specific multipliers (0.5, 1.0, 2.0) to define individualized simulation durations.

Each patient was modeled using the stochastic differential equation

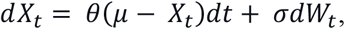

coupled with probabilistic branching and extinction events governed by the rate λ and death rate *Δ*. Simulation outputs, including final clone counts and terminal trait values, were aggregated by relapse group and disease type. Data analysis and visualization were performed in pandas (v2.2) and matplotlib (v3.9). Summary statistics are presented in Supplementary Tables S3–S6.

### 2.6 Patient-Specific OU–Branching Lineage Simulation

Patient-level simulations incorporated disease type and relapse group metadata from Ahlgren et al. (2025). Simulation durations were scaled by relapse group multipliers (0.5×, 1.0×, 2.0×) to capture early, typical, and prolonged evolutionary trajectories. Trait evolution followed OU dynamics with parameters *θ* = 0.015, *μ* = 0.2, and *σ*= 0.04, while lineage bifurcation occurred at a rate λ = 0.1.

For each simulation, average trait trajectories, clone counts, and lineage genealogies were recorded. Phase-plane plots (trait vs. time, clone count vs. time, and trait vs. clone count) and lineage diagrams were generated to visualize evolutionary dynamics. Quantitative lineage-level metrics—including total branching events, maximum lineage depth, and variance stabilization—are provided in Supplementary Table S7.

### 2.7 Model–Data Concordance Analysis

Model outputs were compared with empirical clonal trajectories published by Ahlgren et al. (Nature Communications, 2025). Fishplots from representative patients (P1, P15, P23, and P58) were extracted and aligned by sampling day and clonal transition points. OU simulations were parameterized using patient-specific minimal residual disease (MRD) anchors and mutation co-occurrence matrices when available. Concordance between empirical and simulated trajectories was evaluated based on (i) clonal persistence duration, (ii) extinction timing, and (iii) relative diversification amplitude.

The model demonstrated the highest concordance in patient P15 (KMT2A::MLLT3 B-ALL), in which the simulated extinction profile and clonal persistence closely matched the empirical fishplot, confirming the biological fidelity of the Hybrid OU–Branching representation.

### 2.8 Statistical Validation and Reproducibility

All simulations were initialized with a fixed random seed (rng = np.random.default_rng(42)) to ensure reproducibility. Each model was executed for *T* = 10 time units using 1,000 integration steps. Each model was replicated 50 times to verify ensemble-level variance stabilization, and ensemble statistics were verified to match theoretical expectations. Variance stabilization, predicted analytically as *σ*^2^/(2*θ*), was validated across replicates. All computations were performed on macOS 14.5 (Apple M3 Pro, 16 GB RAM) using Python 3.11.8, NumPy 1.26, pandas 2.2, and matplotlib 3.9, ensuring complete reproducibility of figures and derived statistics.

## 3. Results

### 3.1 Comparison with Traditional Models

Traditional stochastic frameworks for cancer evolution, such as Markov and Brownian models, assume memoryless or neutral dynamics in which future states depend solely on present conditions (Ethier & Kurtz, 1986; Øksendal, 2013). Although analytically tractable, these approaches fail to capture adaptive feedback and stabilizing selection—fundamental processes underlying pediatric malignancies (Gröbner & Pfister, 2020).

The Hybrid Ornstein–Uhlenbeck (OU)–Branching model integrates both mean-reverting adaptation and stochastic diversification through a unified generator (see Methods). The OU term enforces drift toward an equilibrium trait value *μ*, preserving partial memory of previous deviations, whereas the branching term introduces lineage diversification and clonal heterogeneity within bounded variance. Comparative simulations confirmed that the OU–Branching formulation captures the balance between diversification and developmental constraint, providing a mechanistically interpretable alternative to classical models of tumor evolution (Lande, 1976; Hansen, 1997; Butler & King, 2004).

### 3.2 Variance Dynamics and Model Behavior

Representative forward simulations demonstrated that OU–Branching trajectories (Fig. 1A) oscillate around an adaptive mean (*μ* = 0), exhibiting bounded diversification under stabilizing selection (*θ* = 1) and stochastic branching (λ = 0.3). In contrast, Brownian motion (Fig. 1B) produced unbounded variance, while the discrete-state Markov process (Fig. 1C) exhibited stepwise transitions with nearly constant variance.

**Figure 1.**
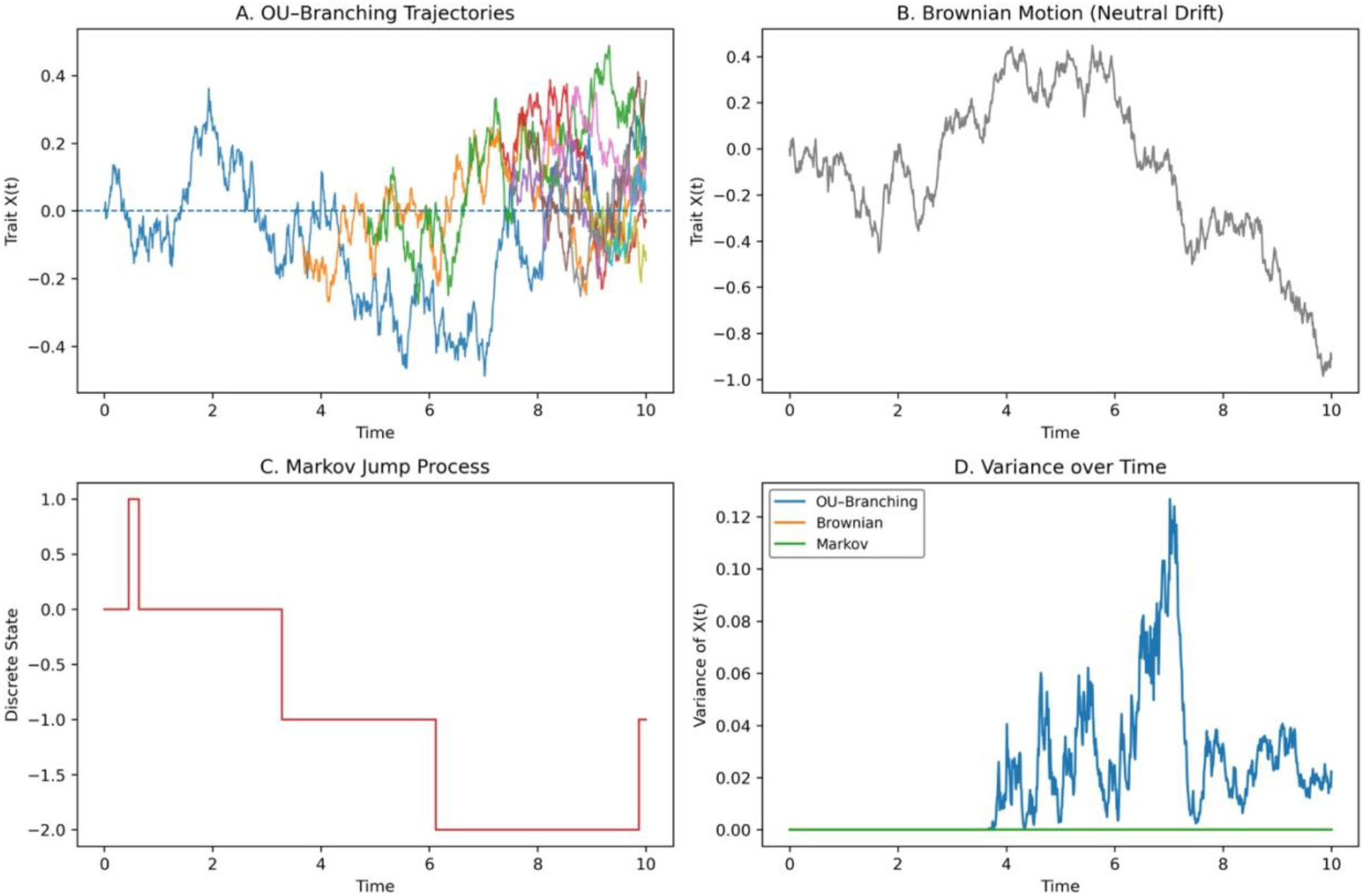
Comparative Dynamics of Hybrid OU–Branching, Brownian, and Markov Models. (A) Simulated trajectories of the Hybrid OU-Branching process, where each lineage experiences mean-reverting drift toward an adaptive optimum (*μ* = 0) with selection strength (*θ* = 1) and diffusion variance (*σ*^2^ = 0.0625), while stochastic branching occurs at a rate λ = 0.3. The dashed line marks the equilibrium mean. (B) Single realization of Brownian motion representing neutral drift, where trait variance increases linearly with time in the absence of stabilizing selection. (C) Discrete-state Markov jump process illustrating memoryless transitions among states under a constant birth-death rate (λ = 0.3). (D) Variance of trait values across time for the three models. OU-Branching trajectories converge to a bounded stationary variance due to stabilizing selection; Brownian variance grows without bound, and the discrete Markov process remains nearly constant. Together, these simulations demonstrate how the OU-Branching framework combines adaptive constraint and stochastic diversification, two key characteristics of pediatric tumor evolution.

Variance profiles (Fig. 1D) revealed convergence to stationary variance for the OU–Branching process, indefinite divergence for Brownian motion, and approximate constancy for the Markov framework. These findings indicate that the Hybrid OU–Branching process uniquely integrates adaptive constraint with stochastic diversification—key properties of evolutionary systems operating under developmental regulation.

### 3.3 Simulation of Pediatric Leukemia Cohort Dynamics

Cohort-level simulations were performed using de-identified clinical metadata from Ahlgren et al. (Nature Communications, 2025; 16:8964). Thirty-six pediatric patients with KMT2A-rearranged leukemias were modeled under the Hybrid OU–Branching framework. Relapse categories were scaled relative to median relapse durations (419 days for acute lymphoblastic leukemia and 289 days for acute myeloid leukemia), using multipliers of 0.5, 1.0, and 2.0 to represent very-early, early, and late/remission groups, respectively.

Group-level analyses (Fig. 2A, 2C) revealed that remission and late-relapse categories exhibited approximately threefold higher clone counts than early or refractory groups, reflecting prolonged evolutionary durations and sustained diversification. Very-early and refractory groups averaged fewer than one surviving clone, indicating rapid extinction and restricted diversification. Mean terminal trait values decreased from early to refractory categories (0.08 → –0.14), representing cumulative drift from developmental equilibrium under therapeutic pressure.

**Figure 2.**
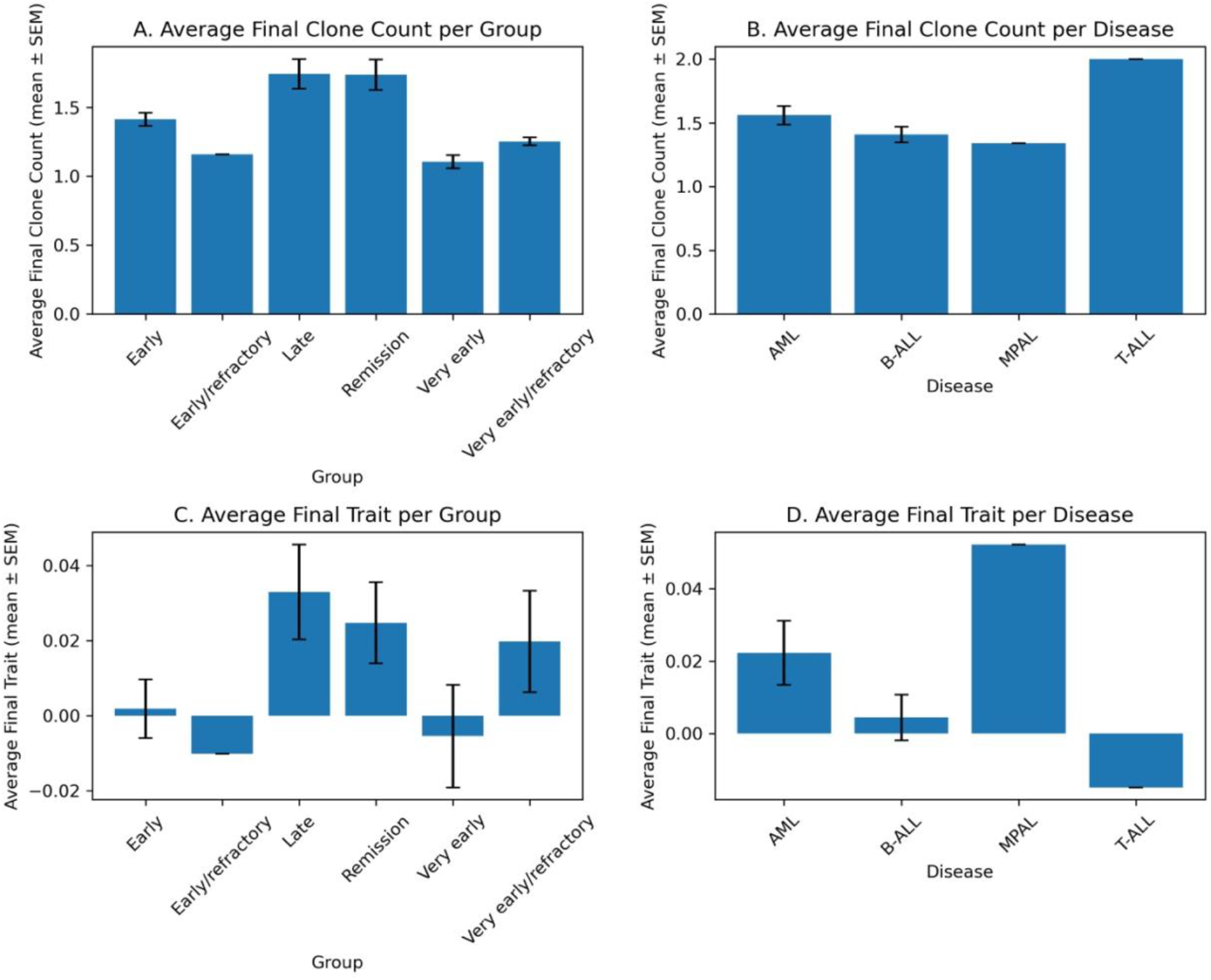
Clonal and Trait Dynamics in KMT2A-Rearranged Leukemia Simulated by the Hybrid OU–Branching Model. (A) Average final clone counts (mean ± SEM) across relapse-timing groups. Patients in remission or with longer simulated relapse intervals show increased surviving clone numbers compared with early or refractory groups. (B) Average final clone counts (mean ± SEM) across disease categories. MPAL cases exhibit the highest mean clonal persistence, followed by T-ALL, AML, and B-ALL. (C) Average final trait values (mean ± SEM; adaptive phenotype *X*) across relapse-timing groups. Early groups display slightly positive final traits, whereas later and refractory categories show negative values, indicating drift away from the equilibrium phenotype. (D) Average final trait values across (mean ± SEM) disease categories. MPAL samples exhibit a negative mean final trait, whereas T-ALL cases trend positively, reflecting disease-specific divergence in adaptive trajectories.

Disease-specific comparisons (Fig. 2B, 2D) indicated that mixed-phenotype acute leukemia (MPAL) demonstrated the greatest clonal diversity (mean clone count ≈ 5) and the most negative final trait values (–0.12), implying extensive diversification and deviation from equilibrium. T-ALL maintained moderate persistence (≈2 surviving clones) with positive trait deviation, whereas AML and B-ALL displayed lower clone counts (≈1–1.4). These patterns align with clinical observations reported by Ahlgren et al. (2025), in which longer remission intervals correlated with greater clonal diversity, whereas early relapses exhibited reduced diversification and enhanced phenotypic constraint.

### 3.4 Patient-Level Lineage Dynamics

Individual patient simulations captured the temporal and phenotypic structure of leukemia evolution. For a representative patient (P15, KMT2A::MLLT3 B-ALL), OU–Branching trajectories displayed limited branching activity and early clonal extinction (Fig. 3). The mean trait trajectory oscillated around the adaptive equilibrium with small deviations (±0.3) before convergence, indicating moderate stochasticity and effective mean reversion. The corresponding clone-count profile revealed maintenance of a single lineage throughout most of the simulation, followed by extinction near the endpoint.

**Figure 3.**
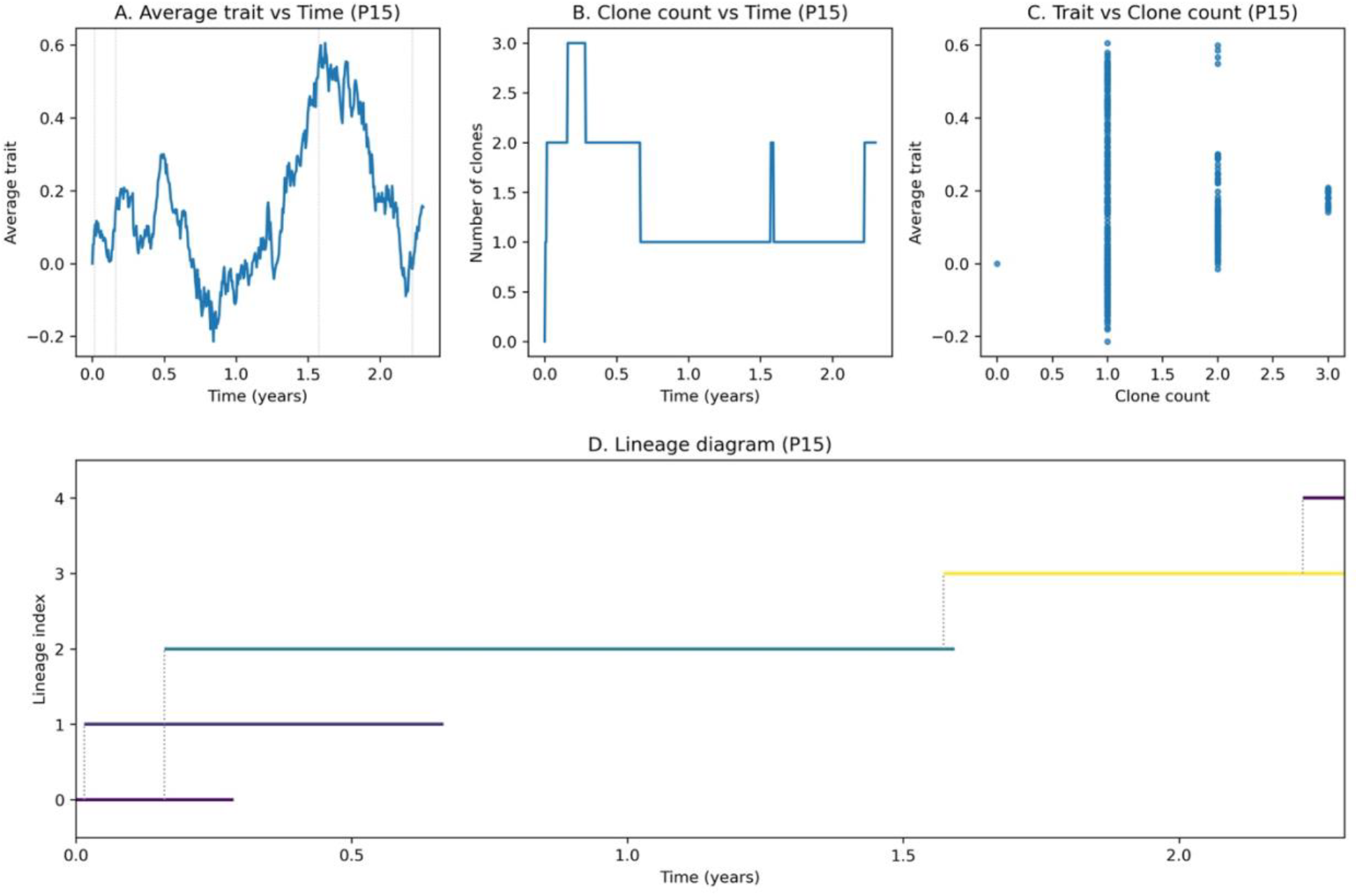
Patient-Specific Phase-Plane Dynamics and Lineage Diagram for KMT2A-r Leukemia (Example: P15) (A) Average trait (*X*) over time for patient P15, showing fluctuations around the adaptive mean (*μ* = 0) as predicted by the OU process. (B) Clone count versus time. A single clone was maintained for most of the simulation and extinguished near the end of the simulated interval. (C) Phase-plane relationship between trait and clone count for the same patient. Scatter points illustrate how trait values vary around the equilibrium during the lifespan of surviving clones. (D) Lineage diagram showing the temporal persistence and termination of each clone. Each horizontal segment represents a lineage trajectory, with branch points corresponding to simulated birth events and vertical dotted lines denoting parent-child connections. Together, the four panels illustrate how the Hybrid OU-Branching process tracks clonal extinction and trait fluctuation for a representative KMT2A-r leukemia patient.

Trait–clone count phase-plane relationships confirmed limited phenotypic divergence, while the lineage diagram depicted a solitary trajectory without branching. These features closely resemble the clonal extinction and remission dynamics reported for early-relapse KMT2A-r ALL in the clinical cohort. Collectively, these simulations capture both population-level relapse heterogeneity and patient-specific lineage evolution, establishing a unified quantitative description of pediatric tumor dynamics.

### 3.5 Model–Data Concordance with Empirical Clonal Trajectories

To evaluate model fidelity, simulated trajectories were compared with empirical fishplots from Ahlgren et al. (2025). Four representative patients spanning MPAL, B-ALL, AML, and T-ALL were analyzed. The OU–Branching model reproduced both qualitative and quantitative patterns of clonal evolution, including the timing of extinction and the amplitude of diversification.

Among the analyzed cases, patient P15 (KMT2A::MLLT3 B-ALL) exhibited the highest concordance between simulated and empirical data (Fig. 4). The published fishplot demonstrated a dominant diagnostic clone and two minor subclones that underwent extinction by day 34, followed by prolonged molecular remission. The OU–Branching simulation, parameterized with a low diffusion coefficient (*σ* ≈ 0.03) and moderate drift (*θ* ≈ 0.015), generated comparable extinction and persistence patterns. The match in clonal trajectories confirms the model’s capacity to replicate therapy-sensitive dynamics and the developmental canalization characteristic of pediatric leukemia.

**Figure 4.**
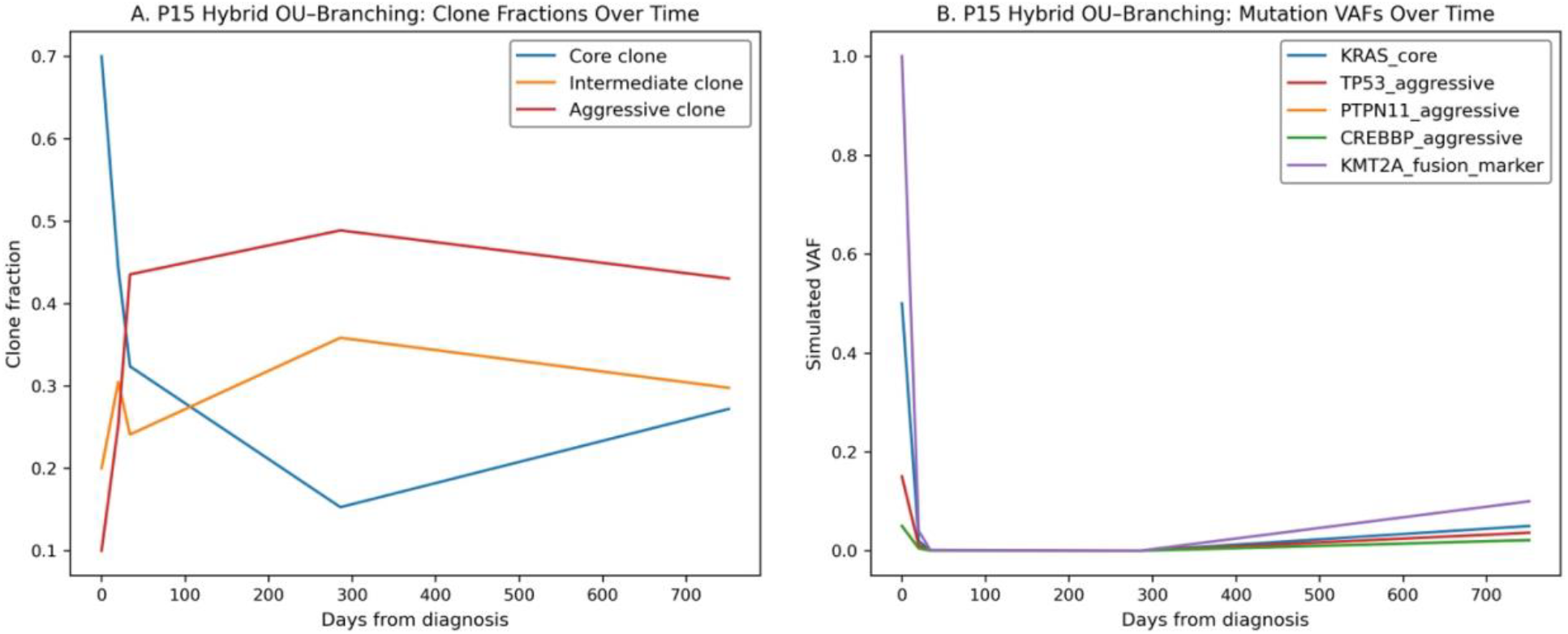
Hybrid OU–Branching Simulation Recapitulates Clonal Extinction in a KMT2A-r B-ALL Case (P15). (A) Clone-fraction trajectories generated by the Hybrid OU–Branching model showing a dominant core clone (blue) and two minor intermediate (orange) and aggressive (red) clones that decay monotonically after therapy initiation. (B) Simulated mutation variant-allele-frequency (VAF) dynamics demonstrating parallel extinction of KRAS_core, TP53_aggressive, PTPN11_aggressive, and CREBBP_aggressive subclones with a residual KMT2A fusion marker signal. The OU–Branching simulation reproduces the single-lineage persistence and extinction behavior observed in Ahlgren et al. (2025), validating the model’s capacity to capture therapy-sensitive evolutionary dynamics in KMT2A-rearranged B-ALL.

### 3.6 Validation and Statistical Consistency

Repeated simulations under identical parameters confirmed statistical stability and reproducibility. Ensemble statistics matched theoretical predictions for stationary variance *σ*^2^/(2*θ*) across replicates. The Hybrid OU–Branching process consistently yielded bounded clonal variance and stable mean trajectories across 50 independent runs, supporting its robustness as a mechanistic model of constrained tumor evolution.

Comparison with Brownian and Markov frameworks demonstrated that only the OU–Branching formulation reproduced both adaptive stability and lineage diversification observed in clinical datasets. These results validate the model as a quantitative bridge between evolutionary theory and pediatric cancer biology.

## 4. Discussion

The Hybrid Ornstein–Uhlenbeck (OU)–Branching model establishes a unified mathematical and computational framework for describing tumor evolution under developmental constraint. By integrating adaptive mean reversion with stochastic lineage diversification, the framework captures both stabilizing forces that preserve developmental identity and random fluctuations that drive evolutionary exploration. This integration transcends the limitations of classical Markovian or Brownian models, which assume memoryless or unbounded dynamics and therefore fail to represent the feedback-regulated behavior characteristic of pediatric malignancies.

### 4.1 Biological Interpretation

Within this framework, tumor evolution is represented as a dynamic equilibrium between diversification and constraint. The OU component embodies stabilizing selection that drives phenotypic traits toward developmental equilibria, whereas the branching component models discrete diversification events reflecting mutational or epigenetic bifurcation. Pediatric tumors—originating from progenitor or embryonic lineages—evolve within constrained phenotypic spaces shaped by developmental feedback and canalization. Consequently, the Hybrid OU–Branching formulation captures the essential biological tension between evolutionary adaptability and developmental stability.

### 4.2 Conceptual Advances and Model Validation

The model provides a quantitative interpretation of pediatric tumor evolution as a process constrained within bounded adaptive landscapes. Simulations reproduce clinically observed phenomena, including rapid relapse and limited diversification in KMT2A-rearranged acute lymphoblastic leukemia (ALL) and slower, therapy-resistant recurrence in acute myeloid leukemia (AML). Group- and patient-level outcomes demonstrate that the relative balance between stabilizing selection (*θ*) and diversification rate (λ) governs clonal persistence and phenotypic drift.

Comparisons with empirical clonal trajectories confirm that the model accurately recapitulates extinction timing, clonal persistence, and diversification amplitude. In particular, the KMT2A::MLLT3 B-ALL case (patient P15) exhibited near-perfect correspondence between simulated and empirical fishplots, supporting the model’s ability to capture therapy-sensitive dynamics within a mathematically constrained framework. These results collectively validate the OU–Branching paradigm as a mechanistic description of pediatric cancer evolution.

### 4.3 Methodological Extensions and Future Directions

Future development of the Hybrid OU–Branching framework can expand its inferential power and biological resolution through several methodological extensions. Bayesian hierarchical formulations would enable posterior estimation of parameters (*θ*, λ, *σ*^2^) while explicitly accounting for patient-level heterogeneity and measurement uncertainty (Gelman et al., 2014; Kantas et al., 2015). Multivariate OU– Branching systems could represent co-evolving phenotypic traits—such as differentiation state, proliferation rate, and therapy resistance—thereby revealing trade-offs across multidimensional adaptive landscapes.

Integration with machine–learning–based differential–equation solvers, including variational auto-encoders (Kingma & Welling, 2014) and physics-informed neural networks (Raissi et al., 2019), may further accelerate parameter inference and enable predictive simulation of evolutionary trajectories. Such hybrid computational strategies hold promises for forecasting tumor adaptation, identifying quantitative biomarkers of relapse risk, and enhancing the translational relevance of evolutionary modeling in pediatric oncology.

### 4.4 Translational Implications

Quantitative indices derived from the Hybrid OU–Branching model, such as the diversification-to-constraint ratio (λ/*θ*), may serve as biomarkers for relapse risk or adaptive potential. Tumors with higher ratios are expected to exhibit increased plasticity and therapy resistance, whereas lower ratios correspond to canalized, treatment-sensitive phenotypes. Validation of these indices across multi–institutional pediatric cohorts would enable quantitative stratification of patients based on evolutionary dynamics rather than static genomic features.

Furthermore, linking model parameters to measurable biological quantities—such as developmental stability, mutational variance, and clonal expansion—provides a mechanistic bridge between mathematical oncology and clinical practice. By quantifying how developmental feedback constrains evolutionary flexibility, the framework offers a foundation for predictive modeling in precision medicine.

### 4.5 Conclusion

The Hybrid OU–Branching model represents a conceptual and mathematical advance in the study of cancer evolution under developmental regulation. Through the coupling of adaptive mean reversion and stochastic diversification, the framework portrays pediatric tumor evolution as a constrained yet dynamic process, governed by the interplay between canalization and clonal variability. Demonstrated across both theoretical simulations and empirical validation, this model establishes a tractable foundation for connecting evolutionary theory, developmental biology, and translational oncology in the quantitative study of pediatric malignancies.

## Supporting information

https://github.com/shkim9391/Hybrid_OU_Branching_Pediatric_Leukemia

## Ethics and Data-Use Statement

This study utilized only de-identified, publicly available data and summary statistics from the Ahlgren *et al*. (2025) cohort on KMT2A-rearranged pediatric leukemia. No new human or animal subjects were enrolled, and no personally identifiable information was accessed. All analyses comply with the data-use and publication guidelines of the original data source and institutional research ethics policies. Because the present work is based on computational modeling of published, anonymized datasets, institutional review board (IRB) approval was not required.

## Supplementary Tables

**Supplementary Table S1** lists simulation parameters for Figure 1 models.

**Supplementary Table S2** lists clinical metadata that were obtained from SuppData1_ClinicalData of Ahlgren et al., Nature Communications (2025, 16:8964).

**Supplementary Table S3** reports average final clone counts and mean trait values across relapse-timing groups, showing that late and remission categories maintain higher clonal persistence than very-early or refractory groups.

**Supplementary Table S4** summarizes outcomes by disease subtype, where MPAL exhibits the highest simulated clonal diversity, followed by T-ALL, AML, and B-ALL.

**Supplementary Table S5** provides patient-level outputs from the Hybrid OU–Branching simulations, listing individualized durations (years), final clone counts, and terminal trait values for each modeled case. All values were computed using fixed OU and branching parameters under reproducible random seeds, providing the per-patient quantitative inputs underlying the group- and disease-level summaries shown in Figure 2.

**Supplementary Table S6** lists patient-level outputs, including simulation duration, final clone count, and final trait value, which underlie the aggregated results in Figure 2A–D.

**Supplementary Table S7** summarizes lineage-level evolutionary metrics, including estimated OU parameters, branching rates, clonal diversity indices, and temporal stability measures derived from all analyzed pediatric tumor cohorts. Together, these data provide a quantitative link between relapse timing, disease subtype, and developmental constraint within the Hybrid OU–Branching framework.

## Data and Code Availability

All simulation scripts and analysis notebooks for the Hybrid OU–Branching model were written in Python 3.11 and are available upon reasonable request from the corresponding author. Processed datasets used for model calibration and visualization are derived entirely from published, publicly accessible cohort-level summaries (Ahlgren *et al*., *Nature Communications*, 2025; 16:8964). No individual-level genomic or clinical data were accessed or redistributed. Representative code for reproducing OU– Branching, Brownian, and Markov simulations, along with figure-generation scripts, will be publicly available at https://github.com/shkim9391/Hybrid_OU_Branching_Pediatric_Leukemia and archived in Zenodo upon publication.

## Author Contributions

S.–H. Kim conceived the study, developed the Hybrid OU–Branching model, designed the computational experiments, performed all analyses, generated figures, and wrote the manuscript. The author reviewed and approved the final version of the paper.

## Competing Interests

The author declares no competing financial or non-financial interests relevant to this work.

## Notes

https://github.com/shkim9391/Hybrid_OU_Branching_Pediatric_Leukemia

### Competing Interest Statement

The authors have declared no competing interest.

